# A Spatiotemporal Atlas of Mouse Gastrulation and Early Organogenesis to Explore Axial Patterning and Project *In Vitro* Models onto *In Vivo* Space

**DOI:** 10.1101/2024.12.12.628063

**Authors:** Luke TG Harland, Tim Lohoff, Noushin Koulena, Nico Pierson, Constantin Pape, Farhan Ameen, Jonathan Griffiths, Bart Theeuwes, Nicola K Wilson, Anna Kreshuk, Wolf Reik, Jennifer Nichols, Long Cai, John C Marioni, Berthold Gottgens, Shila Ghazanfar

## Abstract

At the onset of murine gastrulation, pluripotent epiblast cells migrate through the primitive streak, generating mesodermal and endodermal precursors, while the ectoderm arises from the remaining epiblast. Together, these germ layers establish the body plan, defining major body axes and initiating organogenesis. Although comprehensive single cell transcriptional atlases of dissociated mouse embryos across embryonic stages have provided valuable insights during gastrulation, the spatial context for cell differentiation and tissue patterning remain underexplored. In this study, we employed spatial transcriptomics to measure gene expression in mouse embryos at E6.5 and E7.5 and integrated these datasets with previously published E8.5 spatial transcriptomics^1^ and a scRNA-seq^2^ atlas spanning E6.5 to E9.5. This approach resulted in a comprehensive spatiotemporal atlas, comprising over 150,000 cells with 88 refined cell type annotations as well as genome-wide transcriptional imputation during mouse gastrulation and early organogenesis. The atlas facilitates exploration of gene expression dynamics along anterior-posterior and dorsal-ventral axes at cell type, tissue, and organismal scales, revealing insights into mesodermal fate decisions within the primitive streak. Moreover, we developed a bioinformatics pipeline to project additional scRNA-seq datasets into a spatiotemporal framework and demonstrate its utility by analysing cardiovascular models of gastrulation^3^. To maximise impact, the atlas is publicly accessible via a user-friendly web portal empowering the wider developmental and stem cell biology communities to explore mechanisms of early mouse development in a spatiotemporal context.

## INTRODUCTION

Gastrulation is a critical phase in early embryonic development during which a relatively simple, bi-layered arrangement of cells undergoes rapid proliferation, migration, and differentiation into various mesoderm, endoderm and ectodermal precursors that establish the body plan and give rise to all major organs^4,5^. Although significant progress has been made in profiling these early stages of murine development at both single-cell and molecular levels^2,6–11^, the intricate spatial interactions between cells and tissues as they are specified and patterned during gastrulation remain underexplored. Spatial omics technologies, which allow for the profiling of tissues at the resolution of hundreds of genes, have yet to be applied to the study of these early stages of mouse gastrulation, representing an important gap in our understanding.

We recently generated a comprehensive, time-resolved single-cell RNA sequencing (scRNA-seq) atlas, spanning embryonic (E) days 6.5 to E9.5 of mouse development which encompasses gastrulation and early organogenesis^2^. This atlas, comprising over 400,000 single-cell transcriptomes, defines 88 transcriptionally distinct cell types across embryonic and extraembryonic tissues. Using a previously published cell atlas^6^ spanning E6.5-E8.5, we identified 351 marker genes that distinguish cell types in mouse embryos across this timespan, selected for their suitability in spatial transcriptomics via sequential fluorescent in situ hybridization (seqFISH)^12^. In our previous work, we examined the expression patterns of these 351 markers in sagittal sections of three E8.5 mouse embryos^1^.

In this study, we transform the previous static snapshot spatial data into a temporally resolved spatial atlas of mouse gastrulation and early organogenesis by applying the same seqFISH approach to 20 sagittal sections of four E6.5 and E7.5 mouse embryos. This advanced atlas provides new insights into how heterogeneous transcriptional profiles align with spatial locations throughout mouse development. Additionally, it enables exploration of gene expression dynamics along anterior-posterior and dorsal-ventral axes, shedding light on rapid, region-specific mesodermal fate decisions within the primitive streak at E6.5. Finally, we developed a computational pipeline to project additional scRNA-seq datasets into this spatiotemporal atlas. As a proof of concept, we mapped the anterior-posterior distribution of cell types in mouse gastruloid models^3^, showcasing the atlas as a powerful tool for benchmarking developmental processes *in vitro*.

## RESULTS

### Exploring gene expression patterns during mouse gastrulation using seqFISH

To explore gene expression patterns during murine gastrulation, we performed seqFISH^12^ on 20 sagittal optical sections across four embryos at developmental stages E6.5 and E7.5 (Fig. 1a-e, SFig. 1a). After sample preparation, imaging, cell segmentation, and mRNA dot calling, we computed normalized gene expression levels for 351 genes across 14,794 cells (Fig. 1e). Prior to clustering, we performed Seurat rPCA integration^13^, first between embryos at the same developmental stage (e.g., embryo 1 and embryo 2) and then across stages (E6.5 and E7.5). As a quality control step after the first round of integration, we excluded clusters with abnormal RNA/feature counts, as well as a cluster from the most distal region of the ectoplacental cone in E6.5 embryos (SFig. 1b, red clusters). 16 transcriptionally distinct clusters were identified which we visualized in a low-dimensional UMAP (Uniform Manifold Approximation and Projection) and in their spatial context (Fig. 1f-h and SFig. 1c). Cell types were annotated based on marker gene expression patterns and spatial localization (Fig. 1g-k and SFig. 1c).

**Figure 1:**
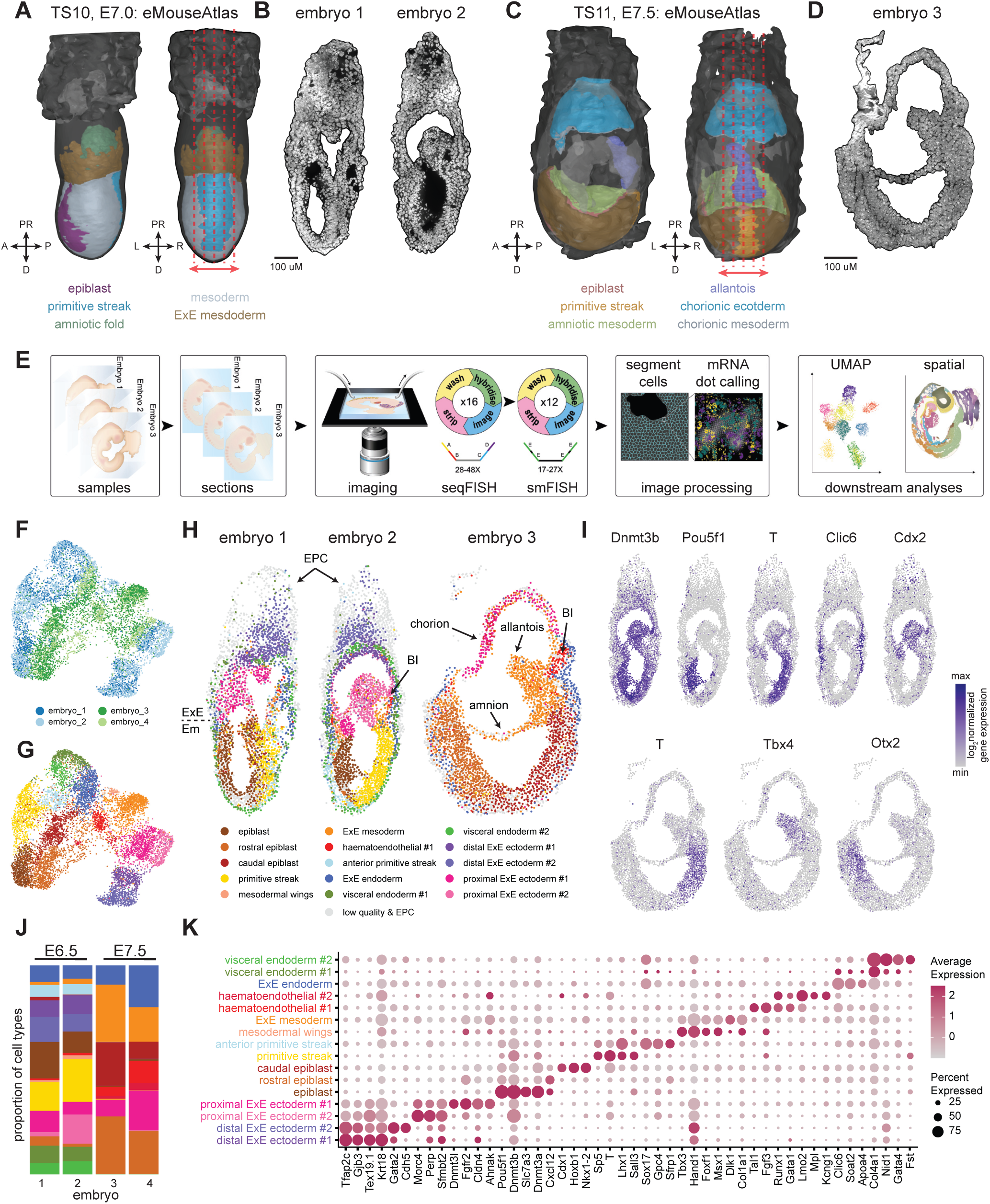
Exploring gene expression patterns during mouse gastrulation using seqFISH. (a, c) 3D illustrations of Theiler stage (TS)10 (a) and TS11 (c) mouse embryos, adapted from eMouseAtlas. Dotted red lines mark the estimated positions of sagittal optical tissue sections shown in (b) and (d). Orientation abbreviations: D, distal; V, ventral; R, right; L, left; A, anterior; P, posterior; PR, proximal. (b, d) Tile scans of 4-μm sagittal sections from two independently sampled E6.5 embryos (b) and one E7.5 embryo (d), imaged using seqFISH with DAPI nuclear staining (white). (e) Schematic outlining the seqFISH pipeline. (f, g) UMAP projections generated from integrated seqFISH expression data. In (f), cells are coloured by their embryo of origin, and in (g), by cell type. (h) Spatial maps of E6.5 and E7.5 embryos, with cells coloured according to their cell types. The black dotted line indicates the extraembryonic (ExE) embryonic (Em) boundary. BI, blood islands; EPC, ectoplacental cone. (i) Representative visualisation of normalised log expression counts of selected genes, measured by seqFISH, to validate performance in E6.5 (top) and E7.5 (bottom) embryos. (j) Stacked bar chart showing the proportion of cell types per seqFISH embryo. (k) Dot plot displaying the average gene expression for marker genes across different cell types identified in the E6.5 and E7.5 seqFISH embryos.

Murine gastrulation begins at E6.25, marked by the formation of the primitive streak at the embryo’s prospective posterior. At this stage, the distal half of the egg cylinder consists of three layers: an outer visceral endoderm, an inner epiblast, and a middle layer of nascent mesoderm and definitive endoderm. The mesodermal layer arises from epiblast cells migrating through the primitive streak region via an epithelial-to-mesenchymal transition. In embryos 1 and 2, we identified an inner epiblast cell layer expressing *Dnmt3a/3b*, *Pou5f1*, and *Slc7a3* (Fig. 1h-k). A prominent posterior primitive streak population expressed *Sp5*, *T*, and *Lhx1*, while anterior primitive streak cells were marked by *Sox17* and *Gpc4* (Fig. 1h-k).

At E6.5, several mesodermal populations were also observed, including extraembryonic (ExE) mesoderm cells expressing *Dlk1* and mesodermal wings at the anterior extraembryonic-embryonic (ExE-Em) boundary in embryo 2, which expressed *Foxf1*, *Hand1*, *Tbx3*, and *Msx1*, and (Fig. 1h,k). Hematoendothelial cells, marked by *Runx1*, *Gata1*, and *Tal1*, were detected in developing yolk sac (YS) blood islands (Fig. 1h,k). Various *Col4a1*+ visceral endoderm populations surrounding the egg cylinder were identified, including an ExE endoderm population expressing *Clic6* and *Soat2* (Fig. 1h-k). Finally, distinct ExE ectodermal populations displayed unique gene expression patterns, in line with their proximal-distal locations relative to the ExE-Em junction (Fig. 1h,k).

By E7.5, the primitive streak has elongated to the distal tip of the egg cylinder as the allantoic bud, arising from the posterior-most region of the streak, begins extending toward the chorionic layer. In seqFISH embryo 3, a prominent allantoic bud composed of ExE mesoderm expressing *Tbx4* was observed (Fig. 1h-k). Additional extraembryonic tissues in embryos 3 and 4 contained ExE mesoderm, including the visceral YS, amnion, and chorion (Fig. 1h and SFig. 1d). E7.5 embryos also contained haematoendothelial cell populations (#1/#2) expressing *Runx1*, *Gata1*, and *Lmo2*, adjacent to ExE endoderm in YS blood islands (Fig. 1h-k and SFig. 1d). In contrast to E6.5 embryos, distal halves of E7.5 embryos consisted of caudal epiblast cells expressing *Cdx1*, *Hoxb1* and *Nkx1-2* and reduced expression of E6.5 epiblast markers (Fig. 1h-k and SFig. 1d) as well as rostral epiblast cells marked by expression of *Cxcl12* (Fig. 1h-k and SFig. 1d).

Altogether, these analyses highlight the utility of seqFISH and our 351-gene probe set to resolve transcriptional diversity associated with spatially arranged cell types at the onset of murine gastrulation.

### Generating an integrated spatiotemporal transcriptional atlas covering murine gastrulation and early organogenesis

To fully leverage these new spatial transcriptomic datasets, we integrated them with three additional E8.5 seqFISH embryo datasets^1^ (SFig. 1d-g) and applied StabMap^14^ and reducedMNN^15^ to align all quality-controlled seqFISH cells (SFig. 1b,d) with a time-resolved scRNA-seq (scRNA) atlas^2^ spanning E6.5 to E9.5 of murine embryogenesis (Fig. 2a,b). After integration, we performed clustering, UMAP visualisation, cell type label transfer, and imputed missing gene expression patterns in the seqFISH embryos. Clusters with poor representation across both datasets, referred to as “poor joint clusters”, were identified prior to further analyses (SFig. 2a). For instance, YS endothelium and YS mesothelium clusters were composed almost entirely (>98%) of scRNA cells, in line with their absence from seqFISH samples. Additionally, seqFISH cells that misaligned with expected embryonic stages (∼12%) were identified and filtered prior to downstream analyses and atlas generation (SFig. 2a).

**Figure 2:**
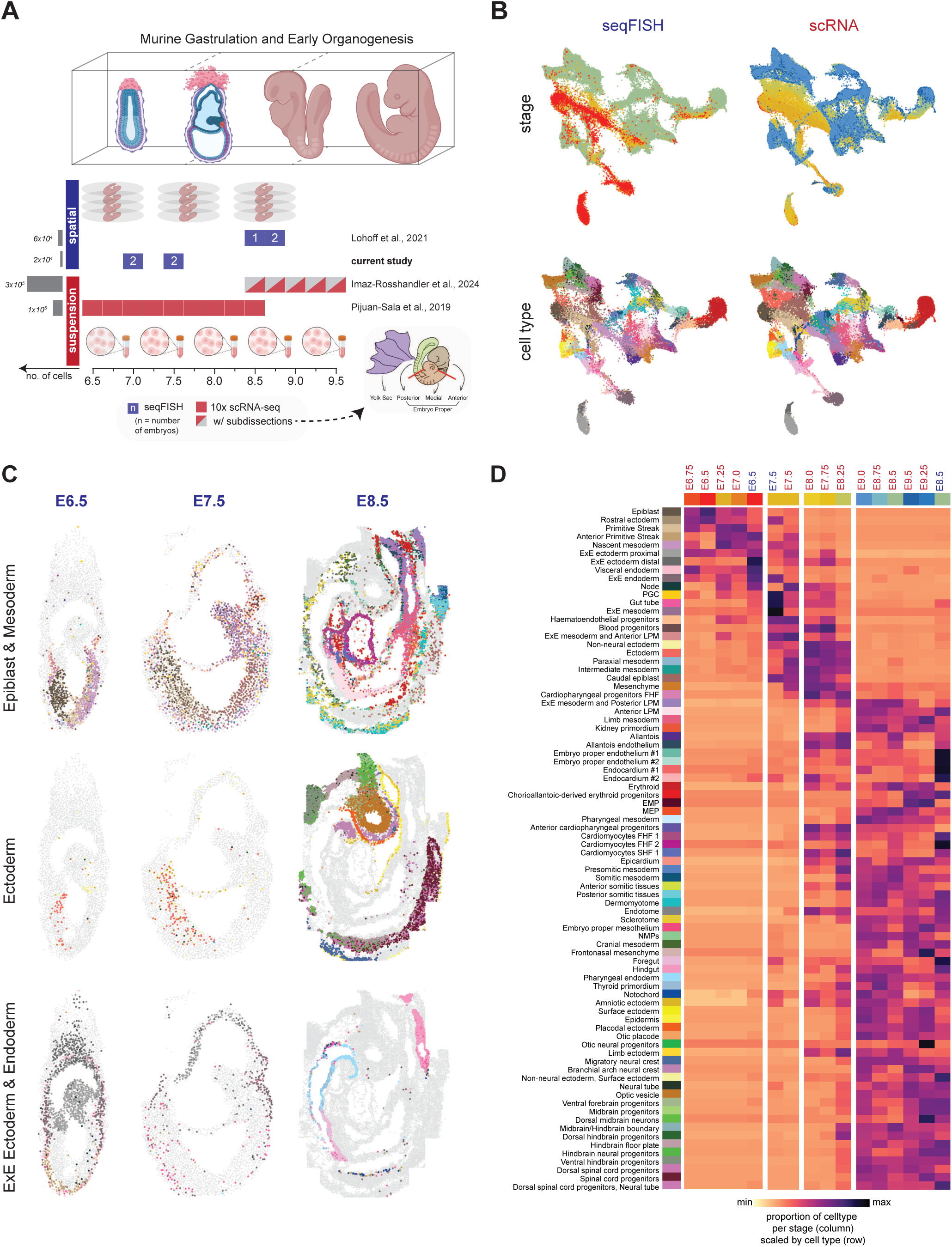
An integrated spatiotemporal transcriptional atlas of mouse gastrulation and early organogenesis. (a) Overview of the single-cell transcriptomic datasets (seqFISH in blue, scRNA-seq in red) integrated to create a spatiotemporal transcriptional atlas of mouse gastrulation and early organogenesis (E6.5-E9.5). Grey bars represent the total number of cells per dataset. (b) Joint projection of cells from the extended gastrulation atlas (scRNA) and seqFISH into a unified reduced dimensional space. Cells are coloured by embryonic stage (top) and refined cell type (bottom). Refer to (d) for the colour legend. (c) Spatial maps of E6.5, E7.5, and E8.5 embryos, organised by germ layer, with cells coloured by refined cell type labels. (d) Heatmap showing the proportion of cell types across embryonic stages in the extended gastrulation atlas (scRNA) and seqFISH samples. Proportions are row-scaled, and the columns were hierarchically clustered.

Cells were initially annotated by assigning joint clusters majority scRNA labels, and these annotations were further refined by analysing marker gene expression patterns and visually inspecting the spatial distribution of cell types within seqFISH embryos (Methods). Altogether, our approach generated a comprehensive spatiotemporal transcriptional atlas of mouse gastrulation and early organogenesis, encompassing 88 refined cell type annotations across more than 150,000 cells (with good alignment from both seqFISH and scRNA datasets) spanning developmental stages E6.5 to E9.5 (Fig. 2b). This new resource enables exploration of cellular diversity in both spatial and temporal contexts during mouse embryogenesis (Fig. 2c and SFig. 2-4), representing the first single-cell resolved spatial map at E6.5-E7.5, and an increase in cell type resolution from 24 to 88 cell types at E8.5.

A comparison of the cellular composition of seqFISH embryos with scRNA embryo pools from different stages revealed E6.5 seqFISH embryos closely align with E6.5–E7.25 scRNA samples, while E7.5 and E8.5 seqFISH embryos correspond to E7.5 and E8.5–E9.5, respectively (Fig. 2d). Detailed counts of cells per cell type across all seqFISH embryos (embryos 1-7) are provided in SFig. 2c-f. Additionally, the spatial distribution of the refined cell types, alongside key marker gene expression patterns from both seqFISH and scRNA cells, are presented by embryonic stage and germ layer in SFig. 2-4. The integrated dataset is accessible through a user-friendly web portal (see Data Availability), allowing researchers to perform virtual dissections, investigate both raw and imputed seqFISH gene expression patterns and identify differentially expressed genes in this context.

### Accounting for spatial location enables refined cell type annotation

Consistent with our analyses in Figure 1, E6.5 and E7.5 seqFISH embryos contained various epiblast-derived cell populations (E6.5–E7.25 scRNA samples), including primitive streak, anterior primitive streak (*Cer1*, *Sox17*), nascent mesoderm (*Mesp1*, *Fgf3*), haematoendothelial progenitors (*Kdr*) and blood progenitors (Fig. 2b-d, SFig. 2g-i). E6.5 and E7.5 seqFISH embryos also contained ectodermal cells expressing *Ptn* and *Cxcl12*, ExE mesodermal cells and primordial germ cells (PGC) (Fig. 2c, SFig. 2g-i). Primitive endoderm-derived populations, including visceral endoderm and ExE endoderm, were identified on the exteriors of egg cylinders, while proximal and distal ExE ectodermal cells localized to extraembryonic regions (Fig. 2c, SFig. 2h). Rostral ectoderm and caudal epiblast cells were present by E7.5. Notably, the higher proportion of distal ExE ectoderm cells in E6.5 seqFISH embryos, compared to scRNA-seq samples, is likely due to the removal of the “sticky” ectoplacental cone prior to sequencing scRNA samples (Fig. 2d).

Previously, we assigned ∼30 cell type labels to E8.5 seqFISH embryos^1^ using a gastrulation atlas covering stages E6.5–E8.5^6^. In this study, aligning E8.5 seqFISH embryos with the expanded gastrulation atlas^2^ (E6.5–E9.5) substantially increased the number of annotated cell types to around 55 (Fig. 2d, SFig. 2b). E8.5 seqFISH embryos contained first and second heart field cardiomyocytes, as well as endocardial cells (#1/#2), cardiopharyngeal progenitors and epicardium which localize to distinctive heart regions in embryos 5–7 (SFig. 3a-f). The allantois endothelium, marked by *Plac1* and *Bambi*, was positioned in the posterior regions of embryos 5 and 7, adjacent to an allantois cell population in embryo 7 (SFig. 3a-f). Embryo proper endothelium (#1/#2), which was depleted in cardiac tissues, contributed to intersomitic and cranial vessels and extended along the trunk (SFig. 3d-f).

Somitic tissues, including the sclerotome (*Pax1*), dermomyotome (*Pax3*), and endotome, were also observed in the trunk, while *Tbx6*, *T, Hes7* and *Dll1/3* expressing somitic and presomitic mesoderm localised to posterior regions, corresponding to the differentiation front—where cells in the presomitic mesoderm transition from a proliferative state to form somites (SFig. 3g-i). The gut tube was composed of pharyngeal endoderm (*Nkx2-3*, *Pitx1*), thyroid primordium, foregut, and hindgut (*Cdx2 and Hoxa7*), which were distributed in distinct patterns along the anterior-to-posterior axes of E8.5 embryos, while notochord (*Noto and Nog*) cells were located dorsally to the gut tube in embryos 5 and 7 (SFig. 4a-c). Additionally, various neural tube and ectodermal cell types were identified in their expected anatomical locations, including spinal cord progenitors, the optic vesicle (*Lhx2*, *Six3* and *Otx2*), placodal ectoderm (*Six3 and Pitx1*) and the otic placode (SFig. 4d-i).

Notably, our spatiotemporal atlas allowed us to refine some of the cell type annotations in the scRNA atlas, including ExE ectoderm and lateral plate mesoderm (LPM) (SFig. 2b). Spatially distinct ExE ectodermal cell types emerge during mouse embryogenesis to form functionally specialised placental lineages. Our spatiotemporal atlas allowed us to identify distal ExE ectoderm cells, expressing *Ascl2* and *Gata2*, located in the ectoplacental cone region at E6.5, which likely include trophoblast stem/progenitor cells^16^ (Fig.2c and SFig. 2g-i). In contrast, proximal ExE ectoderm cells aligned with chorion progenitors, marked by *Elf5* and *Perp*, found adjacent to the epiblast at E6.5 and in the chorion layer at E7.5 (Fig.2c and SFig. 2g-i).

LPM gives rise to a diverse array of critical cell types during murine embryogenesis, including those that form the cardiovascular system, such as the heart and blood vessels^17^. We relabelled a subset of *Hand1*- and *Foxf1*-expressing LPM cells to ExE mesoderm and anterior LPM, as they localised to extraembryonic tissues at E6.5/E7.5 and were present in the medial trunk at E8.5, highlighting transcriptional similarities between ExE mesoderm and intraembryonic medial LPM (SFig. 2g-i and SFig. 3a-c). Additionally, a distinct LPM cluster, marked by *Cdx2* and *Pitx1*, was localized to the posterior of embryos 5–7 and identified in the allantoic bud of embryo 4, prompting its relabelling as ExE mesoderm and Posterior LPM (SFig. 2b, SFig. 3a-c). Overall, our integrated spatiotemporal atlas reveals the spatial and transcriptomic organization of embryonic cell populations, advancing our understanding of mouse gastrulation and early organogenesis.

### Spatiotemporal analysis reveals dynamic gene expression changes along the anterior-posterior and dorsal-ventral axes

During embryogenesis, gene expression patterns coordinate the establishment of embryonic axes, which orchestrate spatiotemporal signals for organ development and delineate the body plan. To investigate gene expression changes along these axes, we assigned normalized anterior-posterior (AP) and dorsal-ventral (DV) coordinates to seqFISH embryos (Fig. 3a and SFig. 5a-g). Next, using our integrated atlas we imputed AP and DV values into the scRNA reference atlas (Fig. 3b and SFig. 5i). Cells from E9.25-E9.5 scRNA samples, isolated from anterior, medial, or posterior regions of sub-dissected embryos, were accurately assigned imputed AP values consistent with their known anatomical origins^2^ (Fig. 3c, see Fig. 2a for sub dissection). This approach provided finer spatial resolution compared to sub-dissection alone, as evidenced by AP gradients within medial and posterior sections (Fig. 3c). Moreover, it enabled the estimation of AP and DV axes in earlier-stage embryos that had not been sub-dissected (pooled, Fig. 3c). The average AP and DV positions of all cell types in seqFISH embryos and the scRNA atlas are shown in SFig. 5h,j.

**Figure 3:**
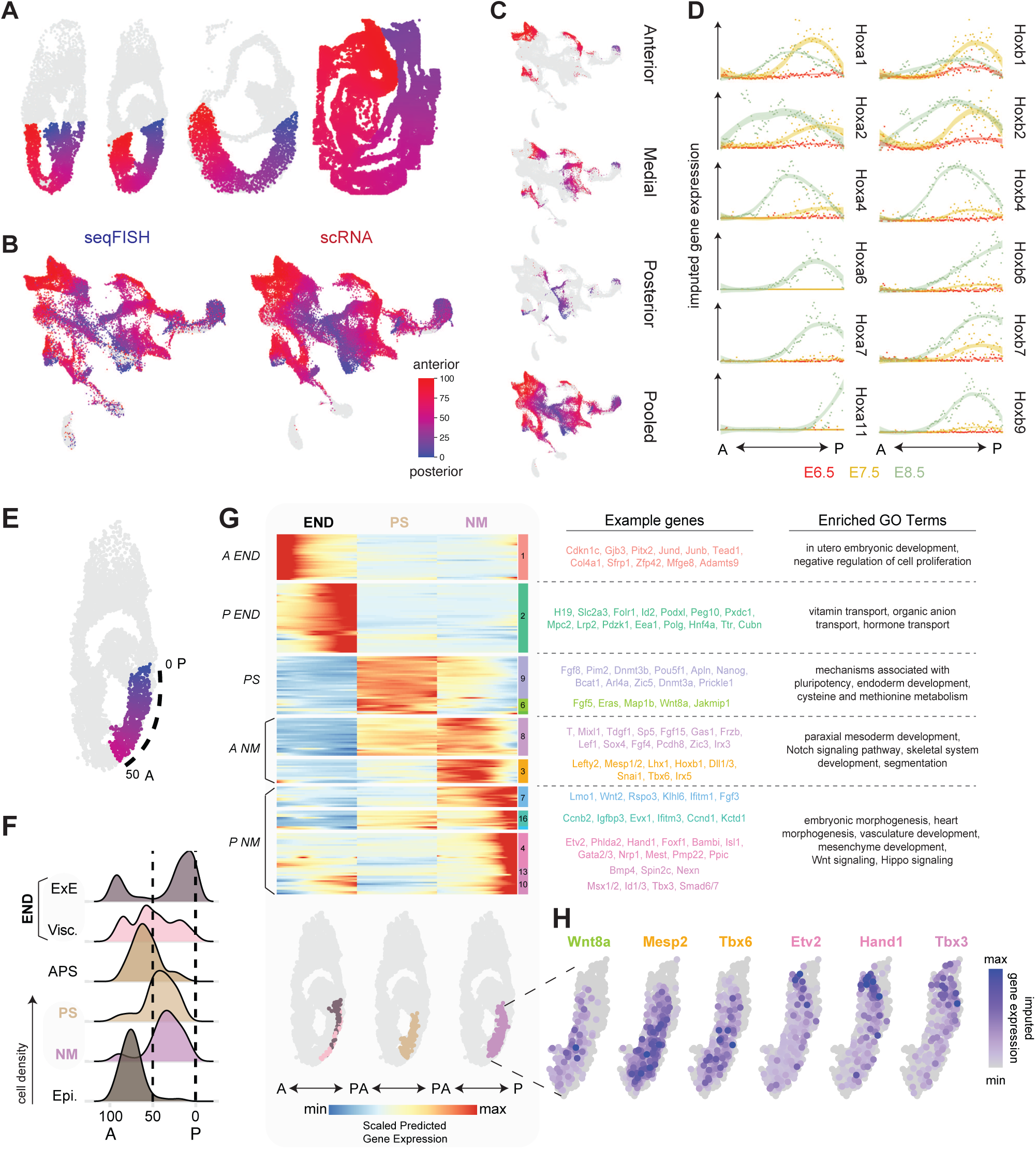
Regionalized gene expression along the anterior-posterior axis highlights rapid fate allocation in the primitive streak during gastrulation. (a) Spatial maps of E6.5, E7.5, and E8.5 embryos, with cells coloured by normalised anterior-posterior (AP) values. Grey cells represent extraembryonic regions or low-quality cells filtered during QC. (b) Joint projection of seqFISH and extended gastrulation atlas (scRNA) cells, coloured by normalised AP values and imputed normalised AP values, respectively (red-blue). (c) Cells from anatomical sub-dissections of the extended gastrulation atlas (scRNA), coloured by imputed AP values. Anterior, medial and posterior sections are from E9.25-9.5 embryos. (d) Binned imputed expression of Hoxa and Hoxb genes along the AP axes in E6.5, E7.5, and E8.5 seqFISH embryos. (e) Spatial plot highlighting primitive streak (PS), nascent mesoderm (NM), and endoderm (END) cells in embryo 2 (red-blue, AP 0-50), with positions indicated by dotted black lines. (f) Ridge plots showing density distribution of cell types along the AP axis in seqFISH embryo 2. (g) Heatmap of smoothed, scaled gene expression profiles along the AP axis in primitive streak, nascent mesoderm and endoderm cells, sorted by AP values and grouped by cell type (END = endoderm , PS = primitive streak, NM = nascent mesoderm). Significant associations identified by Tradeseq^40^ (p < 0.01, mean logFC > 0.25). (h) Visualisation of imputed normalised log expression counts of selected genes, in the primitive streak region indicated in (e).

Incorporating spatial coordinate information into the integrated spatiotemporal atlas provides the developmental biology community with a powerful tool to explore gene expression dynamics along embryonic axes in cell types and tissues of interest at specific timepoints. For instance, we analysed the imputed expression patterns of Hox genes (key developmental regulators with well-established AP-biased expression) along seqFISH embryos from different embryonic stages. Predicted Hox expression patterns (Fig. 3d) align with previous studies^18^, highlighting the utility of our approach.

Next, we investigated gene expression changes in a tissue context along the primitive streak region of embryo 2, focussing on three key populations: primitive streak cells, nascent mesoderm, and the adjacent endoderm layer, spanning positions 0-50 along the AP axis (Fig. 3e,f). This region was selected because mesodermal cells migrating through different AP regions along the primitive streak are known to acquire distinct fates^19–21^. Previous studies have explored bulk RNA-expression patterns in embryonic slices along the AP axis^22,23^. Single cell spatially resolved experiments during this initial process of cell fate diversification, however, have not been performed.

Our analysis of imputed gene expression reveals that numerous genes are dynamically expressed along the AP axis in a germ layer-specific manner within the primitive streak region of embryo 2 (Fig. 3g). Genes associated with the transport of vitamins, organic anions and hormones were enriched in posterior endodermal (P END) cells, while genes related to *in utero* embryonic development such as *Tead1*, *Junb/d* and *Gjb3* were expressed in the anterior endoderm (A END) (Fig. 3g). Primitive streak cells expressed a cluster (9) of pluripotency genes including *Nanog* and *Pou5f1*, that were downregulated in the nascent mesoderm, which displayed limited expression changes along the AP axis. By contrast, the nascent mesoderm expressed gene clusters (3 and 8) in the anterior related to paraxial mesoderm development and Notch signalling, including *Tbx6*, *Mesp2*, *Dll1/3,* while posterior nascent mesoderm was enriched for genes associated with Wnt signalling and vasculature development (clusters 7,16,4,13,10) including *Etv2*, *Hand1*, *Foxf1*, *Isl1*, *Nrp1*, *Msx1/2*, *Tbx3*, and *Wnt2* (Fig. 3g,h).

These findings reveal the nature of rapidly established transcriptional programs along the AP axis in nascent mesodermal cells, distinguishing between posterior ExE mesoderm and anterior paraxial mesoderm, in line with fate biases observed in previous fate mapping studies within the primitive streak. Altogether, this highlights the power of our integrated spatiotemporal atlas to explore gene expression dynamics at single cell resolution across embryonic axes.

### Projecting models of mouse gastrulation into in vivo spatial and temporal contexts

Determining which aspects of embryonic development *in vitro* models faithfully recapitulate is crucial for accurately interpreting experimental outcomes and improving differentiation protocols in stem cell biology. A key strength of our spatiotemporal atlas is its ability to integrate and project external single-cell transcriptional datasets into an *in vivo* spatiotemporal context. To facilitate this, we have developed a bioinformatics pipeline that enables researchers to explore their datasets within the spatial framework of mouse development. As a proof of concept, we projected scRNA-seq datasets from gastruloids, embryonic stem cell-derived models of murine gastrulation, into our atlas (Fig. 4a).

**Figure 4:**
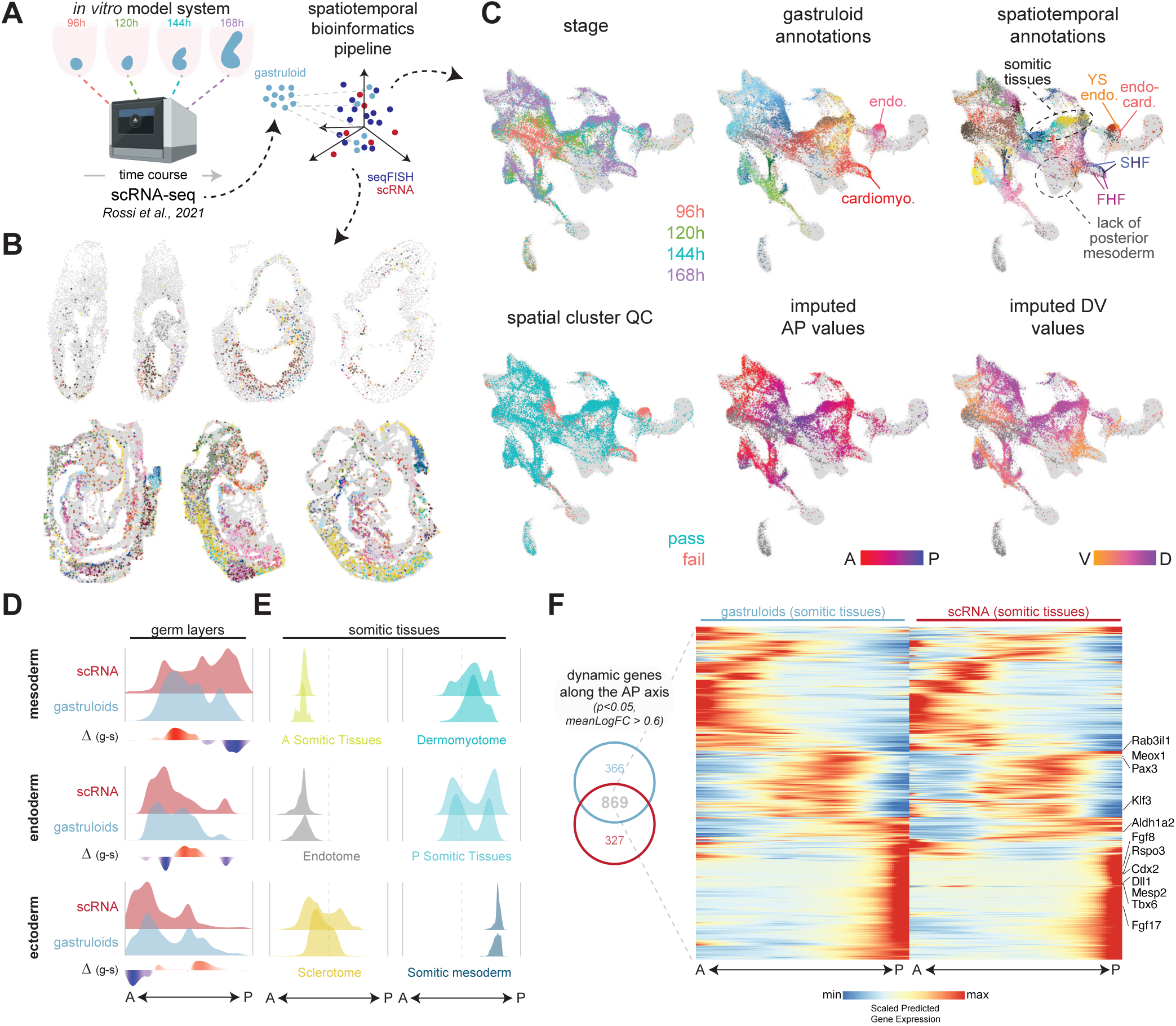
A bioinformatics pipeline to project additional scRNA-seq datasets into a spatial and temporal context during murine embryogenesis. (a) Schematic of the bioinformatics pipeline used to project additional scRNA-seq datasets from from day 4 (96h) -7 (168h) gastruloids, into the spatiotemporal atlas of murine gastrulation and early organogenesis. (b) Spatial maps showing gastruloid cells projected onto the seqFISH embryos in grey, with gastruloid cells coloured according to their cell types. (c) Gastruloid cells projected onto the spatiotemporal atlas, with each panel coloured by different factors. Top row: stage; gastruloid annotations from Rossi et al., 2021; refined cell type annotations from the spatiotemporal atlas. Bottom row: joint cluster quality control (QC); imputed anterior-posterior (AP); imputed dorsal-ventral (DV) values. (d,e): Ridge plots displaying the density distributions of selected cell types from different germ layers (d) or somitic cell types (e) along the anterior-posterior (AP) axis in the scRNA atlas (red) and gastruloids (blue). Density difference plots are shown below (Δ(g-s)), which highlight regions of increased (red) or decreased (blue) gastruloid density relative to the scRNA atlas in (d). (f): Heatmap of smoothed, scaled gene expression profiles along the imputed anterior-posterior (AP) axis for the somitic cell types shown in (e) for gastruloids (left) and the scRNA-seq atlas (right). Significant associations were identified using Tradeseq association test (p < 0.01, mean log fold change > 0.6).

Gastruloids replicate key aspects of anterior-posterior (AP) axis development in the absence of extraembryonic tissues^24–28^. We projected scRNA-seq datasets from day 4 (96h) to 7 (168h) gastruloids, grown using an enhanced induction protocol that promotes cardiovascular development^3^ (Fig. 4a). In line with previous findings^3,29^, our projections highlight formation of first and second heart field cardiovascular progenitors and endocardial cells as well as haematoendothelial progenitors and YS endothelial cells (Fig. 4b,c). It is worth noting that only a subset of gastruloid cardiovascular cells align with seqFISH cells and therefore spatial information is only reliable for those that pass ‘spatial QC’ (Fig. 4c). For example, YS endothelium does not pass spatial QC, so spatial information for this cell type may be less reliable. Beyond cardiovascular progenitors, gastruloids also developed an array of mesodermal, ectodermal, and endodermal cell types (Fig. 4b,c). These included a substantial amount of paraxial mesoderm-derived tissues such as anterior and posterior somites, and other key cell types typically formed during mouse embryogenesis between E6.5 and E9.5 (Fig. 4b,c).

Projection into the spatiotemporal atlas enabled us to assign imputed AP and DV values to the gastruloid cells that passed spatial QC (Fig. 4c). Comparing cell densities of gastruloid versus scRNA-seq cells along imputed AP axes, stratified by germ layer, reveals that this particular cardiovascular gastruloid model does not fully capture the complete range of AP development in mesodermal and ectodermal germ layers (Fig. 4d). More specifically, a subset of the most posterior mesodermal cell types, including posterior lateral plate mesoderm (LPM) and allantois (Fig. 4c), along with the most anterior ectodermal cell types, were underrepresented, highlighting a need for further protocol optimization to better induce these tissues. Nonetheless, these cardiovascular gastruloids successfully generated a diverse array of somitic cell types (Fig. 4c,e).

Taking advantage of the true single cell resolution of our approach, we assessed the predicted AP distribution of somitic cell types and gene expression patterns in gastruloids, induced using the modified cardiovascular protocol, and compared them to *in vivo* somitogenesis using our scRNA-seq datasets (Fig. 4e,f). Previous TOMO-seq studies^25^, which assess gene expression patterns in tissue slices but not at single cell resolution, have revealed that the dynamic transcriptional patterns along the AP axis of individual 120-hour gastruloids, generated using standard somitogenesis-inducing protocols, appear to align with those observed in E8.5 mouse embryos. Our analysis similarly reveals cardiovascular gastruloids generate the full spectrum of somitic cell types typically found along the AP axis in developing embryos (Fig. 4c,e). Moreover, a suite of genes exhibited similar expression gradients along imputed AP axes in both gastruloids and embryonic somitic tissues (Fig. 4f). Many genes showed expression patterns consistent with those previously observed in TOMO-seq-analysed gastruloids^25^, thus (i) validating our approach, (ii) benchmarking the effectiveness of gastruloids in modelling key aspects of early development, and (iii) introducing a broadly relevant strategy for assessing the cellular output of *in vitro* models at true single cell spatiotemporal resolution.

## DISCUSSION

Here, we present an integrated spatiotemporal atlas of mouse gastrulation and early organogenesis, introducing broadly applicable computational strategies to merge spatial and suspension transcriptomic datasets, leveraging the strengths of both technologies. By extracting per-cell coordinates along the anterior-posterior (AP) and dorsal-ventral (DV) axes from spatial transcriptomic data, we accurately predicted the AP and DV positions of cells profiled using dissociation-based single-cell technologies. Additionally, we enriched the spatial transcriptomics dataset by imputing whole-transcriptome gene expression at the single-cell level, creating a comprehensive transcriptional resource for studying mouse embryogenesis within a spatiotemporal framework. To ensure broad accessibility, we developed a user-friendly web portal http://shiny.maths.usyd.edu.au/SpatiotemporalMouseAtlas/, providing researchers with an intuitive platform to explore, visualize, and analyse these complex datasets, and derive new hypotheses to advance our understanding of fundamental biological processes as well as inform new stem cell differentiation strategies for disease modelling and cellular therapy applications.

Our analysis of gene expression changes in the primitive streak of a late-streak stage embryo underscores the utility of assigning axial coordinates. Previous studies that examined genome-wide expression along the AP axis in mini pools derived from tissue slices of mouse gastrulas at similar stages identified gene expression domains associated with mesodermal fate patterning^22,23^. The single-cell resolution of our dataset reveals that these significant transcriptional changes along the AP axis predominantly occur in migratory nascent mesodermal wings rather than in cells immediately exiting the primitive streak. This precise spatial resolution offers fresh insights into the dynamic shifts in gene expression during gastrulation and highlights the potential of our atlas to explore axial patterning in an unbiased manner, providing a powerful tool for understanding gene regulation during embryogenesis.

A recent study^30^ employed whole-mount transcriptomics using cycleHCR to measure the expression of 254 genes in a mid-streak stage mouse gastrula (E6.5) within a 3D context. Consistent with our findings for our earliest timepoint, this single snapshot study identified nine transcriptionally distinct cell clusters, including epiblast, primitive streak, nascent mesoderm, ExE mesoderm, visceral endoderm, and extraembryonic ectoderm, mapped precisely to their spatial locations. As additional spatial transcriptomic datasets emerge, particularly those covering key stages of mouse gastrulation that only measure hundreds of genes, our computational methods can be applied to align these datasets with our spatiotemporal atlas, enabling transcriptome-wide gene expression imputation and refined cell type characterization.

Mapping cardiovascular gastruloids^3^ onto the spatiotemporal atlas using our data integration approach provides a qualitative benchmarking of *in vitro* murine models against the normal spatiotemporal context of mouse development. Our analysis reveals that while these gastruloids effectively recapitulate key aspects of embryogenesis, including anterior-posterior patterning of somitic tissues, they lack specific posterior LPM cell types. Additional datasets from alternative murine culture systems or *in vivo* perturbations, such as drug-treated, mutant, embryoid body or chimera cells, can similarly be mapped and benchmarked using the atlas, although, special care needs to be taken to avoid biases due to the mapping process itself^39^. Moreover, our group and others are actively developing computational pipelines to cross-examine early developmental processes across species^31,32^. Given the sparsity of human spatial reference data during gastrulation, these methodologies will be especially valuable to extend benchmarking practices to emerging human pluripotent stem cell-derived models^33–35^.

Looking ahead, advancing methods to integrate comprehensive spatiotemporal transcriptional atlases with time-resolved 3D embryo models, such as those generated through light-sheet imaging^36^, will pave the way for creating 4D models that predict transcriptome-wide gene expression changes throughout development. Such future approaches will enable detailed exploration of gene expression dynamics across differentiating cellular lineages, providing essential boundary conditions for performing trajectory inference with suspension datasets and offering deeper insights into cell-cell communication during developmental processes. This work will likely contribute to the creation of a ‘virtual tissue’ of mouse gastrulation and early organogenesis, offering an unprecedented tool for modelling and understanding embryonic development at a new level of precision.

### Limitations of the study

Despite the advances we describe in generating this atlas, there are some further avenues to be explored. While our atlas contains spatial coordinates and whole-transcriptome level gene expression, a consistent “average” coordinate system is yet to be built over both spatial and temporal axes. Further work could leverage recent methodological advances in mapping cells between spatial and temporal coordinates that harness optimal transport, such as moscot^37^ or DeST-OT^38^. However, using these methods would only be applicable for spatially resolved cells rather than leveraging the entire set of cells in the spatiotemporal mouse gastrulation atlas.

Moreover, it is important to consider for any abundance-based downstream analysis, cells in the spatial data may be subject to enrichment or depletion biases due to sample selection. Here, we ameliorated this potential issue by comparing cell densities between gastruloids and scRNA-seq-resolved cells rather than with spatial transcriptomic-resolved cells, though it may be that further biases remain across the entire spatiotemporal atlas, in which case further effort should be placed in determining effective matched controls to aid in comparison in the context of a reference atlas^39^.

## SUPPLEMENTARY FIGURE LEGENDS

**Supplementary Figure 1:**
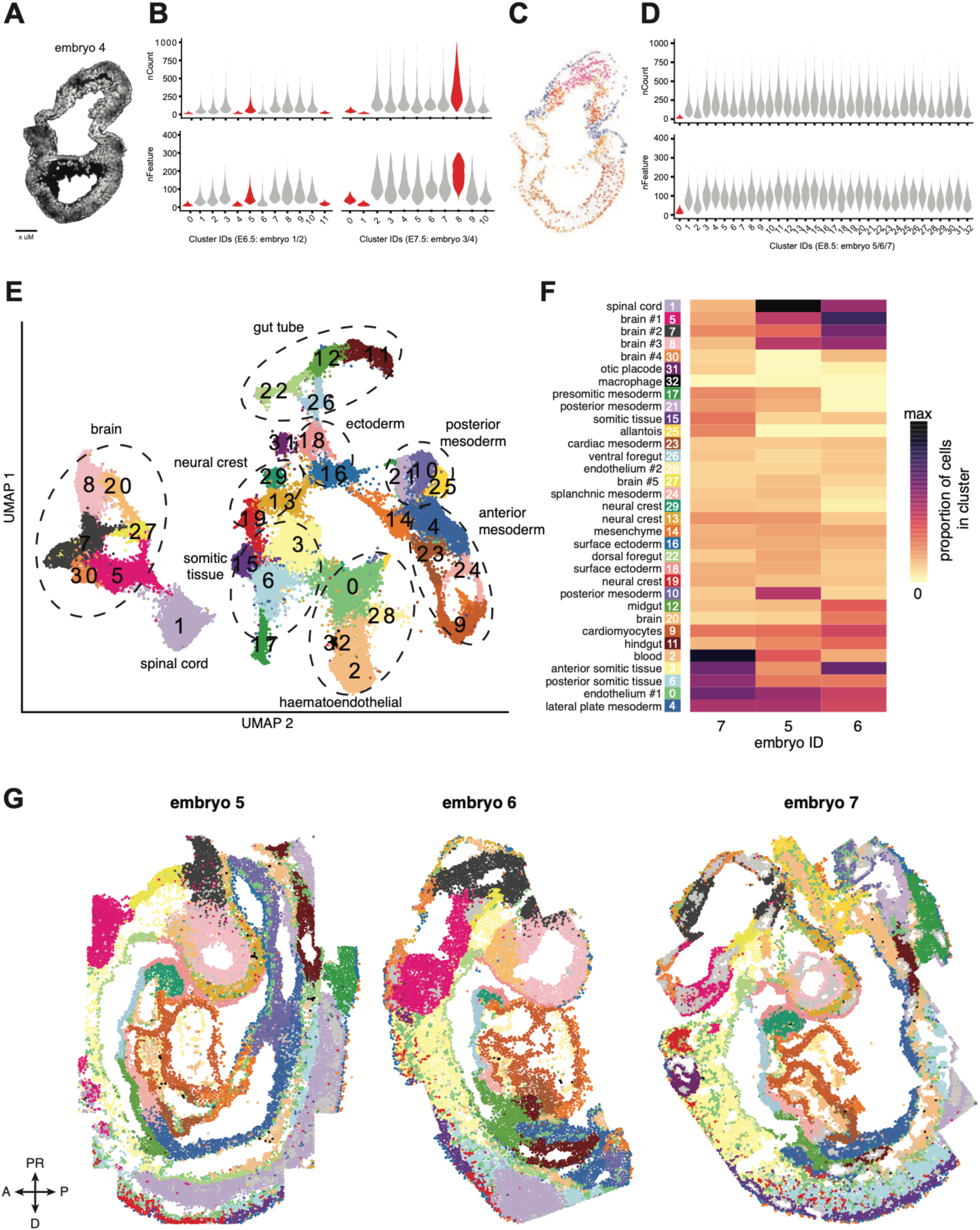
Quality control, clustering, and cell type annotation of E8.5 seqFISH embryos. (a) Tile scan of a 4-μm sagittal section from an E7.5 embryo, imaged using seqFISH and stained with DAPI (white) to highlight nuclei. (b) Violin plots showing the number of counts (top) and number of expressed features (genes, bottom) in cells from unsupervised clusters identified in E6.5/E7.5 seqFISH data. Cells from red clusters (E6.5: 0, 4, 5, 11; E7.5: 0, 1, 8) were excluded from downstream analyses. (c) Spatial map of embryo 4, with cells coloured according to their cell types, as in Fig. 1 h. (d) Violin plots showing the number of counts (top) and number of expressed features (genes, bottom) in cells from unsupervised clusters identified in E8.5 seqFISH data. Cells from red clusters (E8.5: 0) were excluded from downstream analyses. (e) UMAP of integrated E8.5 seqFISH expression data, with cells colored by cell type. (f) Heatmap depicting the proportion of cells per E8.5 embryo assigned to different cell types. Both rows and columns are hierarchically clustered. (g) Spatial maps of E8.5 embryos, coloured by cell type.

**Supplementary Figure 2:**
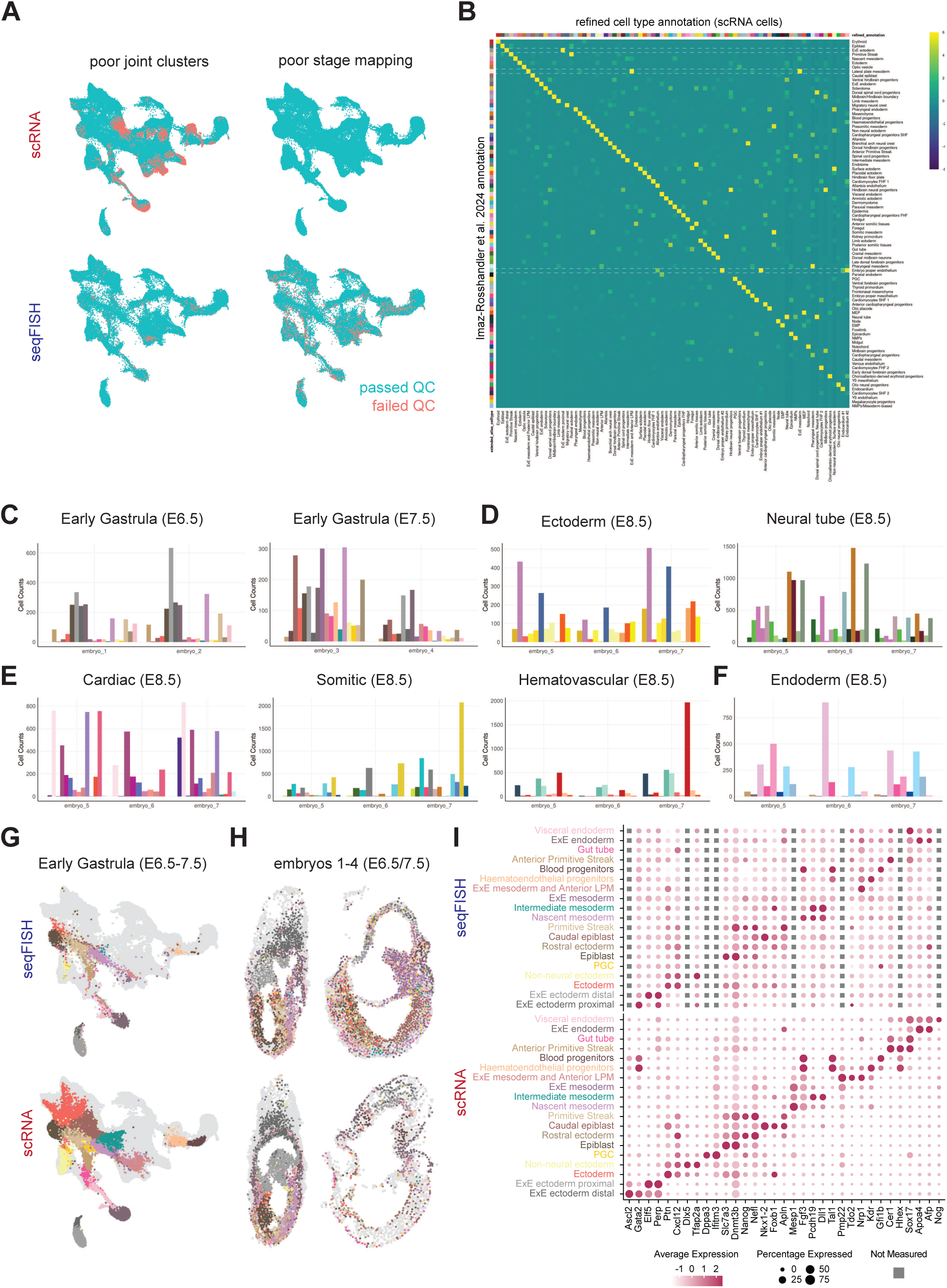
Quality control and refined cell type annotation in the spatiotemporal atlas. (a) Joint projection of cells from the extended gastrulation atlas (scRNA) and seqFISH datasets into a unified reduced-dimensional space. Cells are color-coded based on whether they passed quality control (QC). (b) Confusion matrix showing the number of cells assigned to different refined annotations versus cell types from the extended gastrulation atlas^2^. Rows are scaled, with white lines highlighting ExE ectoderm, lateral plate mesoderm, and embryo proper endothelium, which received further refined cell type annotations. (c-f) Bar charts illustrating the number of cells per cell type across different seqFISH embryos. Bars are coloured according to the refined cell type annotations (see panel b for the legend). A subset of refined cell types visualized in the spatiotemporal atlas (g) or in spatial maps of embryos 1–4 (h). (i) Dot plot displaying the average gene expression of marker genes across different cell types in the seqFISH and scRNA cells shown in (g). Grey boxes indicate genes not measured in seqFISH but identified as marker genes based on imputed gene expression patterns.

**Supplementary Figure 3:**
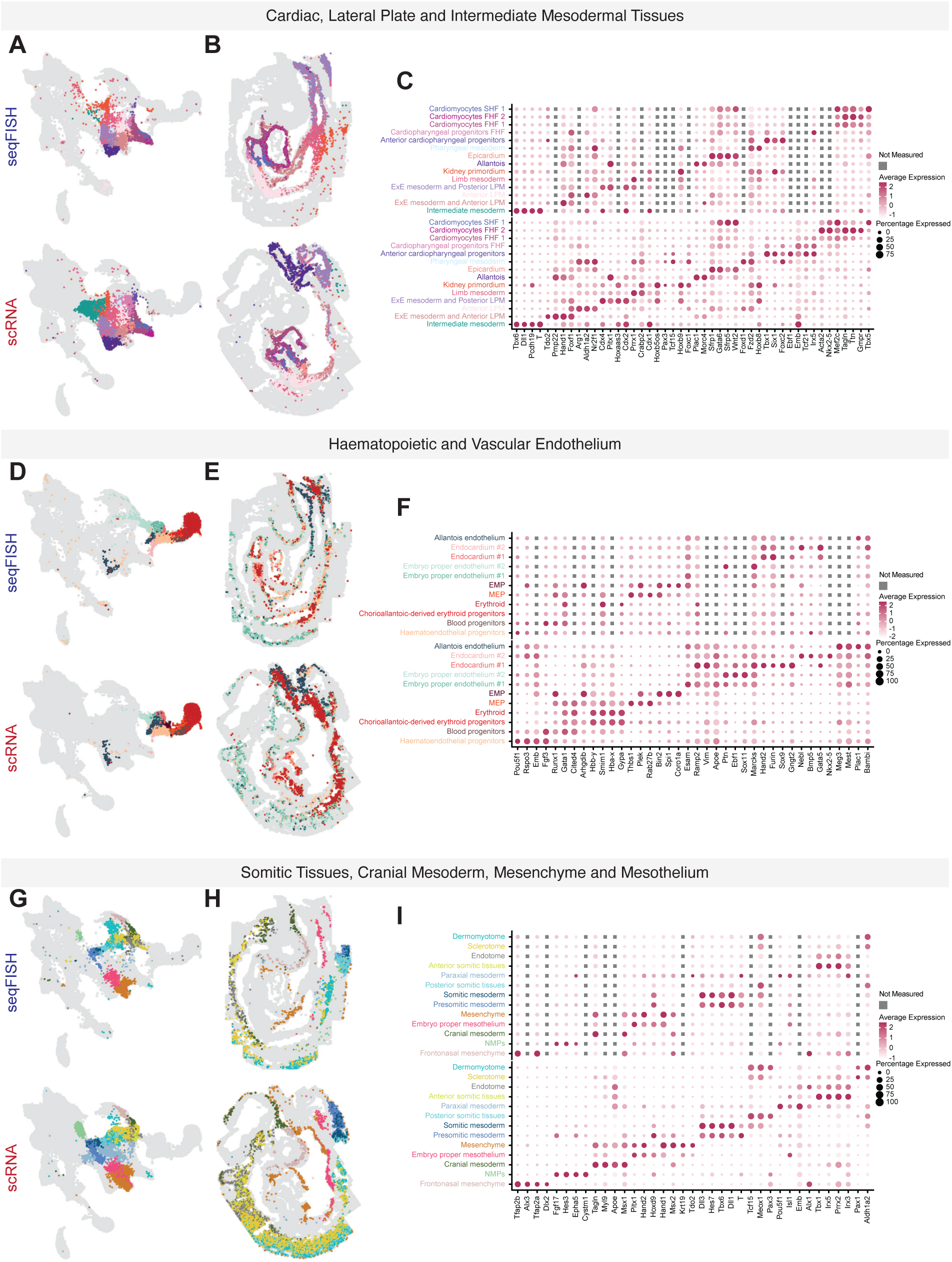
Refined spatiotemporal atlas mesodermal cell types during early organogenesis. A subset of refined cell types visualized in the spatiotemporal atlas (a,d,g) or in spatial maps of embryos 5 and 7 (b,e,h). (c,f,i) Dot plots displaying the average gene expression of marker genes across different cell types in the seqFISH and scRNA cells shown in (a,d,g). Grey boxes indicate genes not measured in seqFISH but identified as marker genes based on imputed gene expression patterns.

**Supplementary Figure 4:**
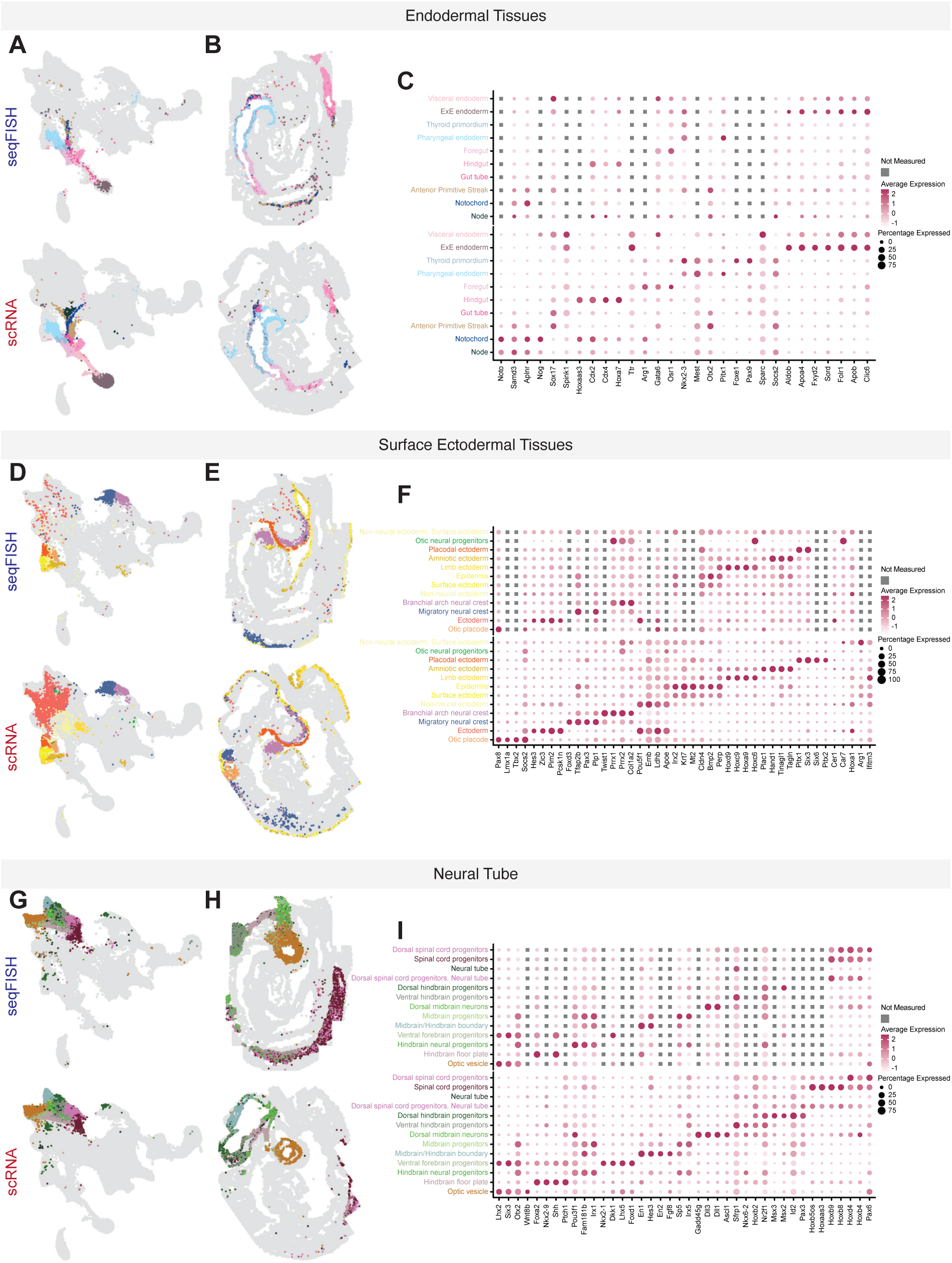
Refined spatiotemporal atlas ectodermal and endodermal cell types during early organogenesis. A subset of refined cell types visualized in the spatiotemporal atlas (a,d,g) or in spatial maps of embryos 5 and 7(b,e,h). (c,f,i) Dot plots displaying the average gene expression of marker genes across different cell types in the seqFISH and scRNA cells shown in (a,d,g). Grey boxes indicate genes not measured in seqFISH but identified as marker genes based on imputed gene expression patterns.

**Supplementary Figure 5:**
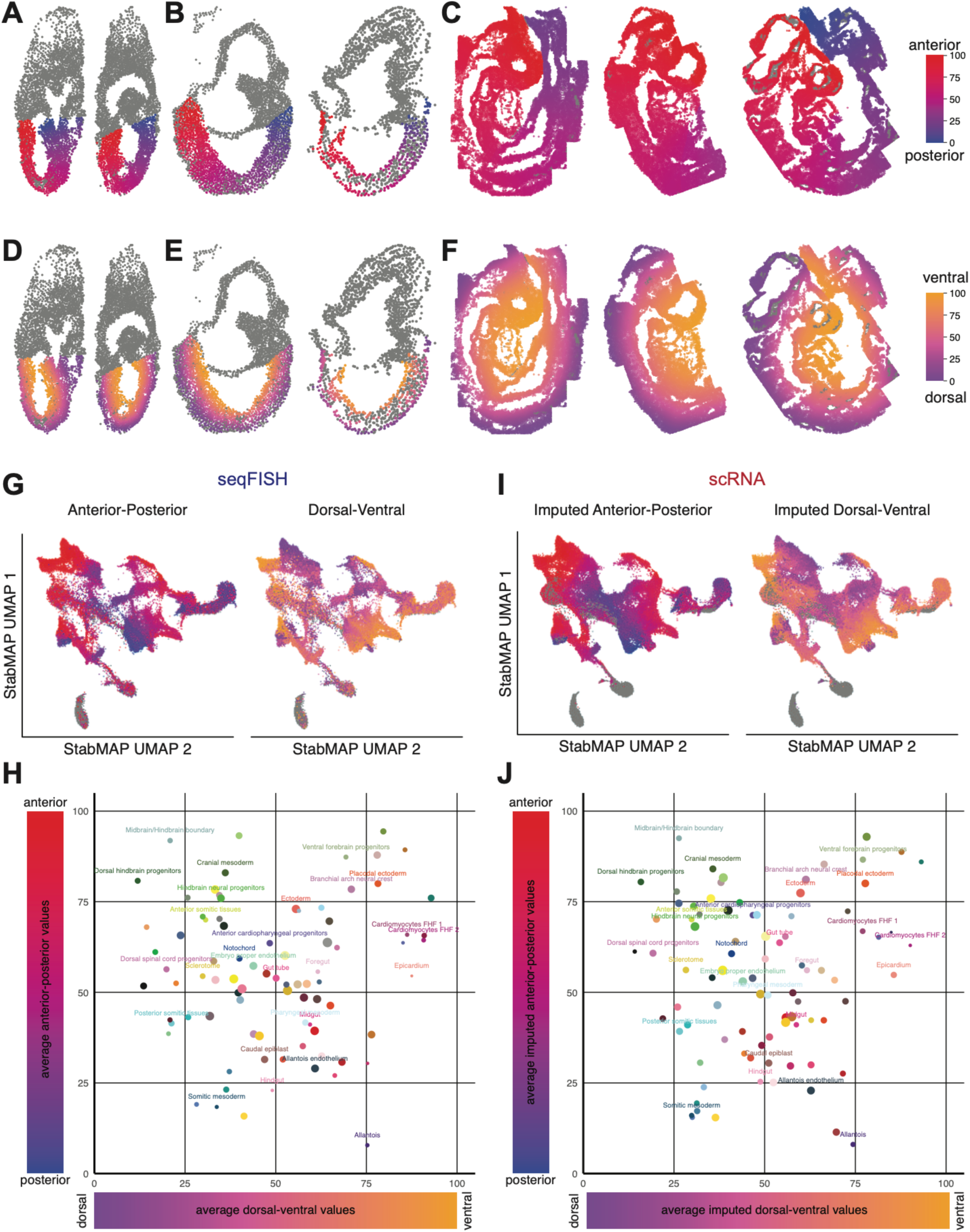
Assigning anterior-posterior and dorsal ventral values to the integrated spatiotemporal transcriptional atlas of mouse gastrulation and organogenesis. (a-f) Normalized anterior-posterior (AP) and dorsal-ventral values were assigned to all cells in the seqFISH embryos. Spatial maps of E6.5, E7.5 and E8.5 embryos, with cells coloured by min max normalized AP values (red-blue, a-c) and percentage rank normalized DV values (orange-purple, d-f). Cells coloured in grey are extraembryonic or are low quality cells that were filtered out during quality control. (g,i) Projection of seqFISH (g) and extended gastrulation atlas cells (i) into the joint reduced dimensional space. seqFISH or scRNA cells are coloured by AP (left) and DV normalized values (right) (h,j) Scatter plot showing the average AP and DV positions of different cell types in the seqFISH dataset (h) and extended gastrulation atlas (j). The size of the dots represents the sum of the variance of the AP and DV values for the cell type.

## MATERIALS AND METHODS

### Library and probe design

We used the same library selection as previously described^1^, using a total of 351 seqFISH barcoded genes and 36 additional non-barcoded sequential smFISH imaging. We used the same primary probe design, readout probe design, primary probe library construction, readout probe synthesis, encoding strategy, and coverslip functionalization as described previously^1^.

### Mice and tissue preparation

Experiments were performed in accordance with EU guidelines for the care and use of laboratory animals, and under the authority of appropriate UK governmental legislation. Mice were used as described previously^1^. Tissue was prepared and sectioned ready for seqFISH imaging as previously described^1^.

### seqFISH microscopy

Three tissue sections from two experimental blocks, containing four embryos, were imaged as previously described^12,41^. For each FOV, snapshots were acquired with 4 um z steps for six z slices. Serial hybridization and imaging were repeated for 29 rounds.

### Imaging processing and registration

Raw image data were processed as described previously^1^. Briefly, effects of chromatic aberration, tissue background and autofluorescence were removed, non-uniform background was corrected and background signal subtracted following the same approach as described previously^1^. For each round of hybridization, Images were registered to the DAPI images of the first hybridization for each FOV using the same two-dimensional phase correlation algorithm as previously described^1^.

### Transcript detection and smFISH processing

We called individual transcript barcodes using the dot matching algorithm with the same parameters as described in Lohoff et al, with more details in Shah et al^41^.For the 36 genes that we probed using smFISH, we found that assignment of an optimal light intensity threshold to separate background noise from hybridized mRNA was particularly challenging due to each gene’s expression only being measured over a single round of hybridization. To address this problem, we extracted a continuous measure of gene expression per cell, taken to be the 95th percentile of the pixel intensities across all pixels within each cell’s segmented area, where a high value corresponds to some evidence of detection of expression of each gene in each cell. We did not use the smFISH-captured genes for further high-dimensional analysis.

### Cell segmentation

Since we visually inspected images associated with the cell membrane channels in the first hybridization and found there was very low signal intensity associated with the cell membranes, we opted for an alternative cell segmentation strategy to what was performed previously^1^. We describe the steps of this segmentation strategy below.

#### Human-in-the-loop cell segmentation

In this situation, we used a human-in-the-loop cell segmentation strategy, where we performed manual annotation with napari (https://zenodo.org/records/8115575) of single points for each DAPI-stained nucleus for each E6.5 and E7.5 image across all z slices. Then, for each z slice and FOV, we performed a seeded watershed algorithm to extend the manually annotated points into solid segments covering the nucleus of each cell. The watershed used nuclear boundary predictions from a U-Net^42^ as heightmap and foreground predictions from the same U-Net as mask. This network was initially trained on the nucleus segmentation dataset^43^ and then retrained several times on the segmentation results obtained via the watershed from manual seed points. Note that we tried using StarDist^44^ and CellPose^45^ for the nuclear segmentation, but found the segmentation quality of their pretrained models insufficient. To further extend nuclei to full cell segments, we trained a feature based classifier to segment the full cell and cell boundaries using ilastik^46^ pixel classification on a few cells using the DAPI channel and the total set of detected transcripts as input channels. We then used a watershed with nuclear segmentations as seeds and ilastik prediction as heightmap and mask to obtain the cellular segmentation.

#### 3D segments for early embryos

Since the z slices were 4um apart, a distance that could conceivably cover a single cell over multiple z slices, we performed a matching algorithm to identify matching 2D segments assigned to a single 3D segment. We applied this matching algorithm to all seqFISH data, including the previously published E8.5 data. The matching algorithm worked by querying consecutive z slices from z slice 2 and so on, identifying for each segment the next segment with the largest overlapping area, and calling the pair of segments a match if the Jaccard index between the pixels was 0.5 or above. Finally, for each matched segment, we calculated the new 3D cell centroid as the centroid of the pixel locations across the x, y, and z planes.

### Data processing

#### Filtering and quality control

Following cell segmentation, we assigned each transcript belonging to the barcoded seqFISH library to the corresponding cell in which the segment belonged. We then filtered cells to retain only those that were detected in i) at least three genes; and contained ii) between 5 and 1,000 total counts. To further check for cells that may be low quality, we performed unsupervised clustering of the log-transformed gene counts, and identified clusters with particularly low total counts to be removed from further analysis.

#### Normalisation

To normalise the gene expression data, we first needed to account for the systematic effect where we observed fewer counts with higher z-slice values. To do this, we first extracted the size factors for embryo 1 and z-slice 2 using the calculation embedded within the ‘logNormCounts()’ function in the scran package, ignoring the Xist gene in the calculation of size factors. We then normalised all of the seqFISH-resolved cells using the ‘logNormCounts()’ function in scran, where size factors were quantile normalised to match the distribution of those of embryo 1 z-slice 2. In doing so, we extracted normalised logcounts for the cells, which we used for subsequent analyses.

#### Defining AP- and DV-axes

To ascertain anterior-posterior (AP) axis positioning of the seqFISH-resolved embryos, using anatomical landmarks, we performed manual annotation of the AP axis, as well as manual annotation of the extraembryonic region and the embryo proper for the E6.5-E7.5 samples. Then we used the ‘princurvè package^47^ to extract the relative positions along the AP axis as well as the perpendicular distances to the AP axis curve to determine the positions along the dorsal-ventral (DV) axis. To enable consistent comparison between each of the seqFISH-resolved embryos, we used z-scaling followed by reordering to start on the anterior part of the AP axis. These scaled AP values were then used for further downstream analysis.

### scRNA-seq Analyses

#### Quality Control Filtering within an Embryonic Stage after Performing Integration and Clustering

Initially, we applied batch correction using the rPCA method in Seurat v4^13^ to log-normalized counts matrices from E6.5, E7.5 and E8.5 seqFISH embryos. rPCA integration was performed across different embryos (e.g., embryo 1 and embryo 2) within each embryonic stage (e.g., E6.5). To perform rPCA integration, principal components (PCs) were generated by running ScaleData and RunPCA using all seqFISH genes set as features. Integration was then carried out using FindIntegrationAnchors (with k.anchor = 5, scale = FALSE, and reduction = ’rpca’), followed by IntegrateData (with features.to.integrate = all features (genes) and k.weight = 100). Finally, clustering was performed after applying ScaleData and RunPCA on the batch-corrected matrices. We used FindNeighbors(dims = 1:30) and FindClusters(resolution = 1) to identify clusters. As a quality control measure, we excluded clusters with abnormal RNA/feature counts, as well as a cluster from the most distal region of the ectoplacental cone in E6.5 embryos (see SFig. 1b,d red clusters) prior to further analyses described below.

#### rPCA Integration, Clustering and UMAP Generation for E6.5 and E7.5 seqFISH Embryos

We applied multiple rounds of batch correction using the rPCA method in Seurat v4^13^ to log-normalized counts from E6.5 and E7.5 seqFISH embryos. Cells that passed the previously described quality control criteria were included in this analysis. In round 1, rPCA integration was performed across different embryos (e.g., embryo 1 and embryo 2) within each embryonic stage (e.g., E6.5). First, principal components (PCs) were generated by running ScaleData and RunPCA on all seqFISH probes. Integration was then carried out using FindIntegrationAnchors (with k.anchor = 5, scale = FALSE, and reduction = ’rpca’), followed by IntegrateData (with features.to.integrate = all features (genes) and k.weight = 100). In round 2, batch-corrected count matrices from round 1 were re-integrated using rPCA, with k.weight = 200 and k.anchor = 5. Finally, clustering was performed on the batch-corrected matrices after applying ScaleData and RunPCA. We used FindNeighbors(dims = 1:30) and FindClusters(resolution = 1.2) to identify clusters, followed by dimensionality reduction using UMAP, generated via RunUMAP(dims = 1:30).

#### rPCA Integration, Clustering and UMAP Generation for E8.5 seqFISH Embryos

For the E8.5 seqFISH embryos, a similar procedure was followed. After quality control, batch-corrected count matrices from within-stage integration were processed with ScaleData and RunPCA. Clustering was then performed using FindNeighbors(dims = 1:30) and FindClusters(resolution = 1), and clusters were visualized via UMAP, generated using RunUMAP(dims = 1:30).

#### Marker Gene Identification

To identify marker genes for the clusters from the E6.5/E7.5 and E8.5 seqFISH embryos, we applied FindAllMarkers(slot = “data”), filtering for marker genes with an average log2 fold change (avg_log2FC) greater than log2(1.5).

#### Calculating Principal Components from the Extended Gastrulation scRNA Atlas for StabMAP

We used log-normalized counts from a downsampled version (10,000 cells per embryonic stage from E6.5 to E9.5) of the mouse gastrulation atlas from Imaz-Rosshandler et al., 2024, to perform three rounds of rPCA integration using Seurat v4^13^. In round 1, rPCA integration was carried out across sequencing batches within each embryonic stage (e.g., E6.5) using 2,000 highly variable features identified with FindVariableFeatures. We applied FindIntegrationAnchors (k.anchor = 5, scale = FALSE), followed by IntegrateData (features.to.integrate = all features (genes), k.weight = 100). In round 2, batch-corrected count matrices from round 1 were integrated across all embryonic stages within each atlas version (original vs. extended). In round 3, integration was performed across both the original and extended atlas versions. For rounds 2 and 3, rPCA integration was executed with k.weight = 200 and k.anchor = 5, using PCs that were identified using a combined list of VariableFeatures (5716 genes) identified from each embryonic stage during round 1 of rPCA integration.

#### Calculating Principal Components from the seqFISH Datasets for StabMAP

We applied multiple rounds of batch correction using the rPCA method in Seurat v4^13^ to log-normalized counts from E6.5, E7.5 and E8.5 seqFISH embryos. Cells that passed the previously described quality control criteria were included in this analysis. In round 1, rPCA integration was performed across different embryos (e.g., embryo 1 and embryo 2) within each embryonic stage (e.g., E6.5). First, principal components (PCs) were generated by running ScaleData and RunPCA on all seqFISH probes. Integration was then carried out using FindIntegrationAnchors (with k.anchor = 5, scale = FALSE, and reduction = ’rpca’), followed by IntegrateData (with features.to.integrate = all features (genes) and k.weight = 100). In round 2, batch-corrected count matrices from round 1 were re-integrated using rPCA, with k.weight = 200 and k.anchor = 5. Finally, principal components were calculated on the batch-corrected matrices after applying ScaleData and RunPCA.

#### Performing StabMAP and reducedMNN to Align seqFISH and scRNA Datasets

To align all seqFISH cells that passed quality control with the downsampled scRNA cells from the extended gastrulation atlas, we applied StabMAP^14^ followed by reducedMNN^15^. For StabMAP, we used the top 30 principal components calculated for both the scRNA and seqFISH datasets, as described above, with projectALL = TRUE. The resulting StabMAP embedding was then reweighted to ensure equal contribution from both datasets using reWeightEmbedding(). To correct for any remaining technical differences between the seqFISH and scRNA datasets, we applied reducedMNN_batchFactor() with k = 10, setting the batch factor to either seqFISH or scRNA as appropriate.

#### UMAP Generation and Joint Clustering for Spatiotemporal Atlas

A low-dimensional UMAP was generated to visualize both scRNA and seqFISH cells within the same space, using runUMAP applied to the batch-corrected StabMAP embedding. Joint clustering and subsequent sub-clustering were performed on this embedding with FindNeighbors (dims = 1:60) and FindClusters (resolution = 1 for clustering; resolution = 3 for sub-clustering). As a quality control step, clusters containing more than 98% or fewer than 2% of either seqFISH or scRNA cells were labeled as ‘poor joint clusters.’

#### Label Transfer across scRNA and seqFISH Datasets

We applied K-nearest neighbors (K = 5) to classify the seqFISH cells based on cell type annotations from the extended mouse atlas, incorporating both extended and original atlas labels, as well as embryonic stage and subdissection labels. Additionally, we used the same K-nearest neighbors approach (K = 5) to classify the scRNA cells according to the cell types assigned to the seqFISH cells.

#### Gene Expression Imputation for seqFISH Cells

We leveraged the scRNA atlas data, which provides whole transcriptome counts, to impute full gene expression profiles for the seqFISH-resolved cells. For each seqFISH-resolved cell, we identified the K-nearest neighbors (K = 5) and calculated the mean expression vector across all genes from the extended mouse atlas dataset.

#### Gene Expression and AP/DV Coordinate Imputation for seqFISH and scRNA Cells

We utilized the seqFISH atlas data, which includes AP and DV axial coordinates, to impute these coordinates in the scRNA cells. To achieve this, we identified the K-nearest neighbors (K = 5) for each scRNA cell and calculated the mean expression vector across the AP or DV coordinates from the seqFISH dataset.

#### Refining cell type annotations using seqFISH data

After obtaining cell type annotations for the seqFISH cells, we visualized their spatial distribution to further distinguish subtypes based on both spatial positioning and gene expression. This analysis was focused on the endothelium, lateral plate mesoderm, and ExE ectoderm. We then back-mapped the scRNA-seq extended mouse atlas to correlate the newly refined cell types with the broader atlas, allowing us to manually annotate subclusters with more specific cell type labels (e.g., proximal and distal ExE ectoderm, anterior and posterior LPM, embryo proper endothelium 1 and 2, and endocardium 1 and 2) based on additional spatial localization information.

#### Marker Gene Identification and Expression Visualization

To identify marker genes for the refined cell types, we used the FindAllMarkers function on seqFISH cells with the imputed gene expression data (slot = “imputed gene expression”) and a curated list of established marker genes from Imaz Rosshandler et al. (2024). This analysis was performed on subsets of cell types (SFig. 2-4), with markers filtered for those exhibiting an average log2 fold change (avg_log2FC) greater than log2(1.5). The original gene expression patterns for these marker genes in both seqFISH and scRNA cells are visualized in dot plots (SFig. 2-4).

#### Differential gene expression analyses along AP axes in the primitive streak region and somitic tissues in the gastruloids and scRNA cells

To identify genes with altered expression over the AP axes in the primitive streak region of embryo 2, or the gastruloids somitic tissues, the AP coordinates were provided as pseusotime values to Tradeseq^40^ (1.10.0; Van den Berge et al., 2020). Generalised additive models were fit (fitGAM, nknots=6, cellWeights=1) to genes that were highly variable [top 2000 highly variable genes were identified using VariableFeatures function in Seurat (4.2.0)] among the cells that were part of each cell type. The associationTest function was then used to identify which of these genes had altered expression over the AP axes. ComplexHeatmap^48^ (2.15.3; Gu et al., 2016) was used to visualise the expression patterns from the GAMs with the highest 300 waldStat scores (*P*-value<0.01 and meanLogFC>2). Metascape^49^ (Zhou et al., 2019) was used to identify Gene Ontology (GO) terms that were enriched for clusters of genes.

### Bioinformatics pipeline to project *in vitro* models into a spatiotemporal framework

To project additional scRNA-seq datasets into the spatiotemporal atlas, we used StabMAP^14^ and reducedMNN^15^. Principal components were calculated for the gastruloid cells after performing rPCA integration using Seurat v4^13^ as described above in the ‘Calculating Principal Components’ sections. Briefly, multiple rounds of rPCA integration were performed, first across samples within each gastruloid timepoint and then across timepoints. Next, to align all gastruloid cells with the spatiotemporal atlas cells, we applied StabMAP followed by reducedMNN. For StabMAP, we used the top 30 principal components calculated for the scRNA, seqFISH and query gastruloid datasets, as described above, with projectALL = TRUE. The resulting StabMAP embedding was then reweighted to ensure equal contribution from all three datasets using reWeightEmbedding(). To correct for any remaining technical differences between the seqFISH, scRNA and gastruloid datasets, we applied reducedMNN_batchFactor() with k = 10, setting the batch factor to either seqFISH, scRNA or gastruloid as appropriate. Cell type labels, and imputed AP and DV values for the gastruloid cells were assigned as described in the ‘Label Transfer across scRNA and seqFISH Datasets” and ‘Gene Expression and AP/DV Coordinate Imputation for seqFISH and scRNA Cells’ sections above.

## ACKNOWLEDGEMENTS

The authors thank all their colleagues, particularly at The University of Sydney, Sydney Precision Data Science Centre, Charles Perkins Centre, and the University of Cambridge Stem Cell Institute for their support and intellectual engagement.

## AUTHOR CONTRIBUTIONS

TL and NK performed the experiments with input from JN, WR, LC. JG and TL performed gene selection with input from JCM and LC. NP performed primary image analysis of raw data with input from LC. CPP and SG performed cell segmentation with input from AK and JCM. LTGH and SG performed analysis of the processed data with input from BT, NW, JCM and BG. FA built the shiny app interface with input from SG. LTGH wrote the manuscript with input from BG and SG. All authors read and approved the final manuscript.

## DATA AND CODE AVAILABILITY

Data can be interactively explored at this link: http://shiny.maths.usyd.edu.au/SpatiotemporalMouseAtlas/

Processed data can be downloaded using the links provided on the front page of the interactive exploration Shiny app above.

All analyses were performed in R (version 4.2.1). Scripts for analysis and figure panels in this manuscript are available at https://github.com/ltgharland/Spatiotemporal-Atlas-of-Mouse-Gastrulation

## Declaration of generative AI and AI-assisted technologies in the writing process

During the preparation of this work the author(s) used ChatGPT in order to improve clarity of writing. After using this tool/service, the author(s) reviewed and edited the content as needed and take(s) full responsibility for the content of the publication.

